# The combined effect of viral infection and temperature on the gene response of melon and zucchini plants with different levels of temperature tolerance

**DOI:** 10.1101/2024.10.18.619003

**Authors:** C. De Moya-Ruiz, M.P. Rabadán, P. Gómez

## Abstract

Biotic and abiotic environmental factors shape plant responses. As such the interplay between viral infection and heat-stress can trigger specific physiological and metabolic plant responses that lead to gene-specific changes in defense and development. However, although plant gene expression patterns have been thoroughly studied under a single stress, the extent to which the combination of both stressors could modulate common or exclusive signaling pathways remains unclear. In this study, we examined the effects of watermelon mosaic virus infection and diurnal temperature variations (20/14 °C, 26/20 °C, and 32/24 °C) on the gene responses of two plant species (melon and zucchini), each with high- and low-temperature tolerance, using a differential 3’mRNA-seq approach. The WMV load was much greater in zucchini than in melon plants, and was also dependent on the temperature conditions and tolerance of each plant species. Our comparative RNA-seq analysis revealed that the percentage of differentially expressed genes (DEGs) was higher in the thermo-susceptible plants of both species under the combination of WMV infection and low temperatures (20 °C). Among these significantly regulated genes, between 37 % and 45 % were related to biotic and/or abiotic stress. Furthermore, we found that 30 GO terms were involved in the response to both combined stress from low temperatures and 23 GO terms for high temperatures, which were exclusive to the thermotolerant varieties. Together, these findings allowed the identification of two unique orthologous genes linked to temperature and virus infection in melon and zucchini plants. Understanding the effects of biotic and abiotic factors on plant responses is essential for unraveling the complexity of plant-pathogen-environment interactions and developing strategies to enhance plant resilience and productivity under changing climatic conditions.

## INTRODUCTION

Plants must adapt to a myriad of abiotic and biotic stresses to thrive. Extreme temperatures, drought, salinity, and greenhouse gases can limit plant growth by triggering specific physiological processes and metabolic responses that lead to gene-specific changes in plant development (Mittler *et al*., 2012; Bailey-Serres *et al*., 2019; Zhang *et al*., 2022). Consequently, climate change poses a significant threat to global food production by affecting agricultural yields, food quality, and prices (Vermeulen *et al*., 2012; Raza *et al*., 2019; Thiault *et al*., 2019). As climate change intensifies, crops also face the impact of pests and pathogens, which could further limit food production and exacerbate food insecurity (Oerke, 2005; Pandey *et al*., 2017; Rudgers *et al*., 2020; Montes & Pagán, 2022; Vasquez *et al*., 2022; Xiao, 2022; Tonnang *et al*., 2022; Laine, 2023). Thus, understanding the intricate interactions between abiotic and biotic stressors is essential to mitigate the potential consequences on agricultural food production systems and further adapt crop management practices to protect food production (Shelake *et al*., 2024).

Heat stress can cause crop production losses by directly interfering with plant physiological processes and reproduction, increasing photorespiration and transpiration rates, altering pollen viability and fertilization, and disrupting metabolic processes (Zhang *et al*., 2022). This effect has been observed in several crops such as beans, cowpeas, wheat, and tomatoes (Bita & Geräts, 2013; Grossiord *et al*., 2020; Karki *et al*., 2021; Rangaswamy *et al*., 2021). Plants can be particularly sensitive to high temperatures during their reproductive stages, which can influence gene expression at the chromatin level and circadian clock (Chang *et al*., 2014; Mody *et al*., 2020), and alter their profiling in plant roots (Martins *et al*., 2017). Moreover, it has been reported that transcriptional factors, including heat shock proteins, are key in regulating gene expression networks involved in plant heat stress responses (Li *et al*., 2019; Tolosa & Zhengbin, 2020; Pardo-Hernández *et al*., 2024), as well as second messengers during signal transduction, protein folding, and protein-protein interactions (Zhang *et al*., 2022). Conversely, suboptimal temperatures can reduce enzymatic activity and biochemical reactions, thereby adversely affecting plant growth and development (Hasdai *et al*., 2006). Furthermore, temperature can modulate plant defense responses through specific proteins, such as NB-LRR proteins (Zhu *et al*., 2010). Understanding how temperature modulates gene expression in plants can allow the identification of genes and alleles that are useful for marker-assisted selection, which can help plant breeding programs to enhance crop resilience (Hill & Li, 2022; Shen *et al*., 2022). Nevertheless, the effect of these improvements on the development of heat-tolerant varieties may be largely contingent on different agroecosystems and pathogen infections (Savary *et al*., 2019).

In addition to plant growth and development, rising temperatures can indirectly influence crop yield by altering plant disease progression and insect pest biology (Jones, 2016; Jeger *et al*., 2018; Jones & Naidu, 2019; Trebicki, 2020; Tonnang *et al*., 2022). Accumulating evidence suggests that crops are vulnerable to viral diseases at high temperatures (Tsai *et al*., 2022b,a; Iqbal *et al*., 2023). In this sense, it has been reported that temperature can influence plant-virus interactions, affecting symptom expression and virus accumulation during the plant infection process (Aguilar *et al*., 2015; Obrępalska-Stęplowska *et al*., 2015; Chung *et al*., 2016). For instance, studies have suggested that virus accumulation is temperature-dependent, with seasonality affecting virus-plant interactions and virus dynamics during persistent infections (Honjo *et al*., 2020), thus, shaping viral genetic diversity and population dynamics in mixed infections (Alcaide *et al*., 2021; Sardanyés *et al*., 2022). Other studies have shown that exposure to elevated temperatures may either enhance or reduce plant susceptibility to viral diseases (Prasch & Sonnewald, 2013; Ghandi *et al*., 2016; Makarova *et al*., 2018; Amari *et al*., 2021; Tsai *et al*., 2022a). However, temperature can also affect the distribution of vectors that transmit plant viruses, leading to more rapid and widespread dissemination (Jones, 2016; Islam *et al*., 2020; Trebicki, 2020). Hence, rising temperatures could accelerate epidemic development and hinder disease management (Jones & Naidu, 2019). However, the interaction between temperature and plant viral diseases is complex and involves changes in vector behavior, viral replication rates, and plant defense mechanisms, among several other biological aspects. This underlines the importance of understanding the molecular mechanisms behind temperature-mediated changes in gene expression, which may play an important role in plant responses to viral diseases. Despite evidence that temperature and viral infections are critical factors that elicit specific responses in plants, their combined effects on gene expression related to defense and developmental mechanisms remain unclear, as they have been extensively studied individually.

To elucidate the interactive molecular responses to combined temperature stress and virus infection, we examined the gene expression profiles of two important cucurbit crops (melon and zucchini) with high- and low-temperature tolerance, and in combination with the presence or absence of the same virus infection, watermelon mosaic virus (WMV). Cucurbits are important horticultural crops in temperate conditions, and are facing increased challenges from viral infections due to the expansion of cucurbit crops, along with changes in cultural practices, and globalization (Grumet *et al*., 2021). Temperature stress (cold and heat), including drought and salt stress, can negatively affect melon growth and productivity (Nguyen *et al*., 2024). Functional genomic studies on plant crops are usually challenging because of the lack of full genome information. Thus, only a few studies have reported specific genes involved in the regulation of the drought stress response in melon plants (Xing *et al*., 2020; Yang *et al*., 2020), and studies on genes involved in heat stress in cucurbit plants are limited. Additionally, 28 viruses have been reported to significantly affect cucurbit crops in the Mediterranean Basin (Lecoq & Desbiez, 2012; Radouane *et al*., 2021). Among these, aphid-transmitted viruses are widely distributed and particularly detrimental to crop production sustainability (Lecoq & Katis, 2014; Moya-Ruiz *et al*., 2023; Rabadán *et al*., 2023). In this scenario, *Potyvirus citrulli* (watermelon mosaic virus, WMV), is a vector-borne virus primarily transmitted by several species of aphids, causing serious diseases in major cucurbit production areas globally (Rabadán & Gómez, 2023; De Moya-Ruiz *et al*., 2021; Rabadán *et al*., 2023). Symptoms of WMV in cucurbit plants include mosaics, chlorotic ringspots, mottling spots, leaf deformation, and plant stunting (Moya-Ruiz *et al*., 2023). Here, we comprehensively integrated and analyzed the viral accumulation and transcriptomic datasets of melon and zucchini plants with high- and low-temperature tolerance, subjected to three temperature ranges (low, medium, and high) and WMV infection (Fig. 1). After analyzing viral RNA accumulation, we conducted a comparative transcriptomic (3’mRNAseq) analysis of single and combined stress responses, characterizing gene clusters with distinct transcription patterns associated with abiotic (temperature) and biotic (viral infection) stress. Additionally, we linked the gene profiles of key biological processes to temperature- and virus-responsive pathways in melon and zucchini plants, which could help develop effective strategies to improve cucurbit productivity.

**Figure 1.**
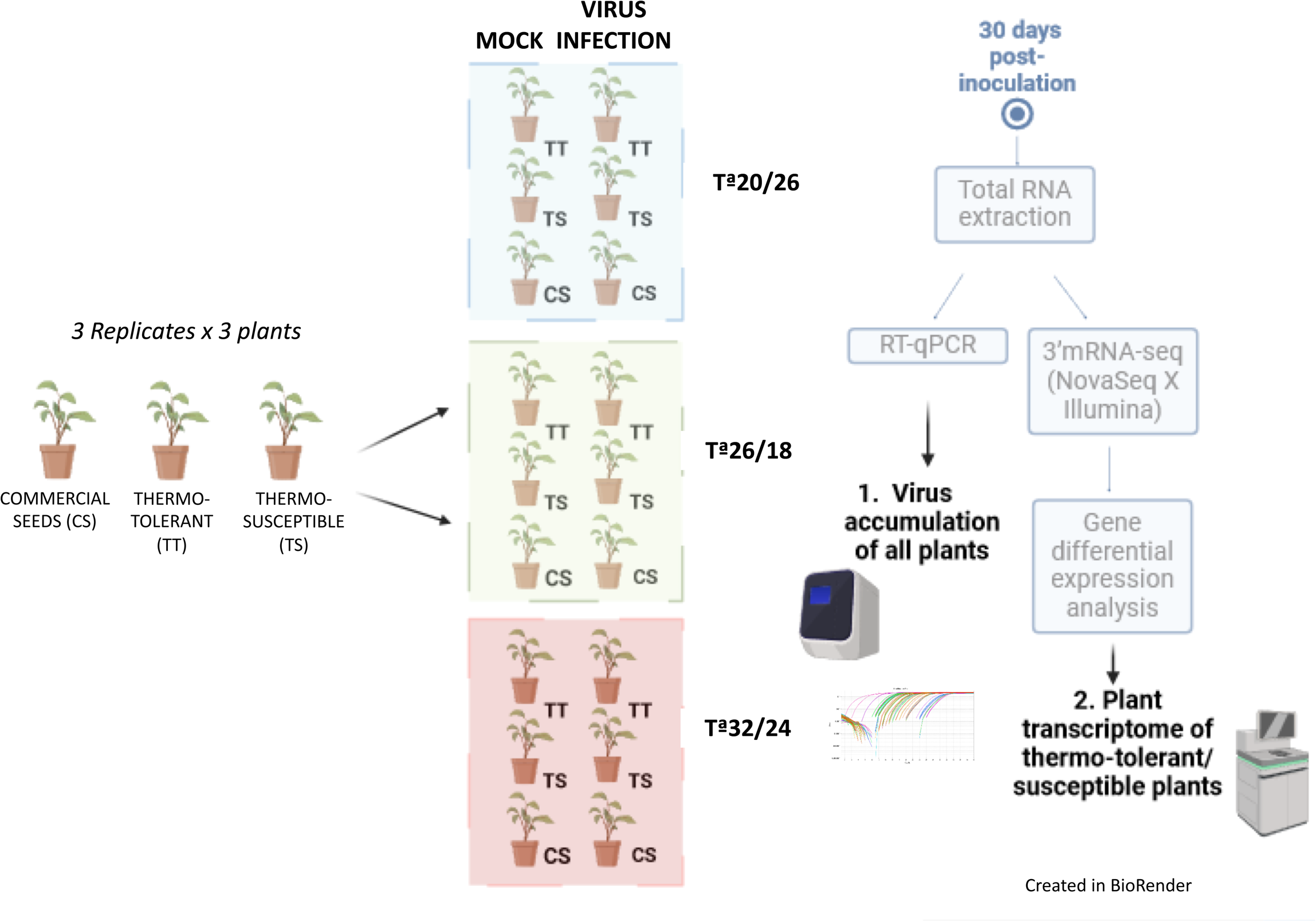
Schematic representation of experimental design and workflow using thermo-tolerant/susceptible melon and zucchini plants. Three replicates, each consisting of a pool of three plants, were maintained at low (20/14 °C), medium (26/20 °C) and high (32/26 °C) temperature ranges under both uninfected and WMV-infected conditions. Plant material was collected at 30 dpi, and WMV accumulation was inferred by absolute quantitative RTq-PCR. Additionally, differential gene expression analysis was conducted using 3’mRNA-seq approach via Illumina (NovaSeq X).

## MATERIALS & METHODS

### Plant material and growth conditions

Each experiment was carried out independently in a controlled greenhouse with a 16/8 photoperiod (16h light and 8h dark) at the respective experimental temperatures: 20°C/16°C (Low), 26°C/20°C (Medium) or 32°C/24°C (High), with ± 6°C of variation between experiment and also day and night condition. Despite the temperature and light regulations, note that the 26°C experiment was conducted from April to June, then the 32°C experiment from July to September, and finally the 20°C experiment from October to December, coinciding with the average temperatures in our place (Region of Murcia, Spain). A total of 108 plants were used for the experiments. This included 9 plants each of thermo-tolerant (TT) and thermo-susceptible (TS) of melon and zucchini varieties (3 replicates x 3 plants). Due to confidentiality agreements and potential conflicts of interest, the specific seed details provided by the respective plant breeding companies cannot be disclosed. Additionally, 54 commercial plants (9 plants each of melon (Piel de sapo) and zucchini (Black beauty) varieties were grown under the same conditions. As a control, mock plants were similarly subject to temperature stress without viral infection (Fig. 1).

### WMV virion purification and inoculation

Plants were inoculated by using WMV virion particles. Briefly, first *Nicotiana benthamiana* plants were agroinoculated with WMV-MeWM7 (De Moya-Ruiz *et al*., 2021) and maintained in a growth chamber at 24 °C. After 20 dpi, approximately 60 g of symptomatic leaves were collected and ground in extraction buffer with liquid nitrogen, following a series of centrifuges and a PEG precipitation as described in (Rupar *et al*., 2013) with some modifications. First, the homogenate was mixed with 5.38 ml/g of homogenization solution (0.5M K_2_HPO_4_, 5 mM EDTA, 10 mM DIECA and Na_2_SO_3_) and agitated at 4°C for 15 min. The mixtures were centrifuged at 7000 rpm for 10 min at 4°C. The supernatant was collected and filtered through gauze, and Triton X-100 was added to the final concentration of 1% and shaken at 4°C for 1h. The supernatant was ultracentrifuged at 50 000 rpm for 90 min at 4°C. After discarding the supernatant, the pellet was resuspended in 14 ml of citrate solution pH 7.5+1% triton X-100 and shaken for 30 min at 4°C. Once again, the pellet was resuspended in 6 ml citrate solution pH 7.5+1% triton X-100, shaken for 30 min at 4°C, and centrifuged at 10 000 rpm for 10 min. The supernatant was collected and 10% chloroform was added, mixed thoroughly, and centrifuged at 10 000 rpm at 4°C for 10 min. The aqueous phase was collected and ultracentrifuged with a 30% sucrose cushion at 45 000 rpm for 2 h at 4°C. The pellet was resuspended in 1 mL of citrate solution and kept overnight at 4°C. The purified virus was stored at −20°C. For plant inoculation with WMV, the third and fourth true leaves were rubbed with Carborundum-dusted and a suspension of 100 mg/mL virions particles in sodium phosphate buffer (30mM), as previously described (Gomez et al., 2009). The material (all apical leaves from each plant) was collected at 30 dpi from 9 plants (3 replicates x 3 plants) per treatment.

### Quantification of viral RNA accumulation

To determine the WMV accumulation, samples were collected at three different time points: 10, 20, and 30 days post-infection (dpi). Total RNA was extracted from all samples using Tri-reagent, purified by phenol-chloroform extraction and treated with DNaseI (Sigma-Aldrich, St. Louis, USA). Viral accumulation was then quantified by absolute real-time quantitative PCR (qPCR) with an AB7500 System (Applied Biosystems, Foster City, CA), using the One-step NZYSpeedy RT-qPCR Green kit, ROX plus (NZYTech, Lisboa, Portugal). Two specific primers targeting the WMV P1 region (523–635 nt) were used: CE-2959 Fw *5’- CACCCAACCTCTGAAATGG-3’* and CE-2960 Rv *5’-GGCTCAGATTTGCATC-3’*. The reaction mix was prepared according to the manufacturer’s instructions (NZYTech), and non-template controls were included to ensure product-specific amplification and the absence of primer-dimers. Serial dilutions (10-fold) of viral RNAs from WMV-MeWM7 infectious clone were used to generate external standard curves (De Moya-Ruiz *et al*., 2021). Initial RNA concentration was measured twice with a Qubit 3.0 fluorometer following the manufacturer’s instructions (Thermo Fisher Scientific). RNA concentration in each sample (ng of viral RNA per 100 ng of total RNA) was estimated by plotting the threshold cycle (CT) values from each biological assay (n=9, at each time-point) with three experimental replicates for each biological replicate. Given that the viral load of WMV in melon and zucchini plants had a similar exponential pattern, consistent with our previous studies (De Moya-Ruiz *et al*., 2021), and most significant differences between temperatures were observed after 30 dpi, subsequent analyses were focused on samples from this time point to ensure the reliability of the results and to capture the relevant biological effects of each temperature condition.

### 3’mRNA sequencing and comparative analysis of gene expression profilling

Total RNA was extracted from samples of melon and zucchini at 30 dpi using Tri-reagent, purified by phenol-chloroform extraction and treated with DNaseI (Sigma-Aldrich, St. Louis, USA). The quantity and quality of RNA were assessed using a NanoDrop ND- 1000 spectrophotometer (Thermo Fischer Scientific, Waltham, MA, USA) and agarose gel electrophoresis. The 3’mRNA-sequencing was performed in a NovaSeq X (Illumina Platform) by Seqplexing (Paterna, Valencia). This RNA-seq approach involves tagging the 3’end of mRNA poly(A) tails with universal adapters, barcodes, and unique molecular identifiers, allowing for accurate quantification of gene expression profiling (Charpentier *et al*., 2021). Briefly, the bioinformatics pipeline starts with cleaning raw sequence data by quality trimming and removing sequencing adapters and poly-A sequences. Quality assessment using FastQC ensures high-quality FASTQ files. UMIs are identified using umi-tools, duplicates are removed and reads are mapped to the reference genome using STAR. The version of the genomic reference for zucchini and melon used were “Cpepo_genome_v4.1”, with the associated transcript annotation being “Cpepo_4.1”, both files obtained from http://cucurbitgenomics.org. and “Harukei3_v1.41” with the associated transcript annotation being “Harukei3_v1.41”, both files obtained from https://melonet-db.dna.affrc.go.jp/ap/top, respectively.

HTSeq-count and counts were normalized to the total number of identified reads in each sample. This inter-sample normalization was followed by control-sample normalization, including control samples (i.e. mock plants under the same stress condition) for each dataset to enable comprehensive comparisons of the gene expression profile between thermotolerant and thermo-susceptible melon and zucchini cultivars. The comparisons, along with the number of Differentially Expressed Genes (DEGs) and total genes from both melon and zucchini cultivars, are summarized in Table 1. DESeq2 was employed for identifying significant expression differences, identifying genes with adjusted p-value < 0.05 and log_2_FC <-1 or >1 for each variety and crop. The variability between groups was analyzed using Principal Component Analysis (PCA) conducted using iDEP 2.0 (http://bioinformatics.sdstate.edu/idep/) (Ge *et al*., 2018). Upset plots and Venn Diagrams were created using ChiPlot (https://www.chiplot.online/) and R package, respectively.

**Table 1.** Summary of the 3’ mRNA-seq experiment design and results of DEGs for each condition including up and downregulated genes under the p-value < 0.05 and Log_2_FC < −1 o > 1 criteria in melon and zucchini varieties.

| Crop | Comparative | Total DEGs/Total gene | %DEGs | DEGs (p-value < 0.05 / Log2FC < -1 o > 1) |  |
| --- | --- | --- | --- | --- | --- |
|  |  |  |  | Upregulated genes | Downregulated genes |
| *Cucumis melo* | TT Melon MOCK Ta20 vs TT Melon WMV-Ta20 | 14955/37254 | 40.14 | 337 | 119 |
|  | TT Melon MOCK Ta26 vs TT Melon WMV-Ta26 | 14882/37254 | 39.94 | 115 | 220 |
|  | TT Melon MOCK Ta32 vs TT Melon WMV-Ta32 | 15351/37254 | 41.20 | 519 | 1102 |
|  | TS Melon MOCK Ta20 vs TS Melon WMV-Ta20 | 15150/37254 | 40.66 | 118 | 110 |
|  | TS Melon MOCK Ta26 vs TS Melon WMV-Ta26 | 10643/37253 | 28.56 | 50 | 115 |
|  | TS Melon MOCK Ta32 vs TS Melon WMV-Ta32 | 10544/37253 | 28.30 | 133 | 280 |
|  | TT Melon MOCK Ta20 vs TT Melon MOCK Ta26 | 15819/15819 | 42.46 | 763 | 682 |
|  | TT Melon MOCK Ta32 vs TT Melon MOCK Ta26 | 15299/15299 | 41.06 | 758 | 477 |
|  | TS Melon MOCK Ta20 vs TS Melon MOCK Ta26 | 15478/15478 | 41.54 | 1126 | 897 |
|  | TS Melon MOCK Ta32 vs TS Melon MOCK Ta26 | 15029/15029 | 40.34 | 820 | 390 |
| *Cucurbita pepo* | TS Zucchini MOCK Ta20 vs TS Zucchini WMV Ta20 | 21938/27868 | 78.72 | 109 | 299 |
|  | TS Zucchini MOCK Ta26 vs TS Zucchini WMV Ta26 | 15963/27868 | 58.28 | 183 | 295 |
|  | TS Zucchini MOCK Ta32 vs TS Zucchini WMV Ta32 | 16281/27868 | 58.42 | 366 | 640 |
|  | TT Zucchini MOCK Ta20 vs TT Zucchini WMV Ta20 | 10825/27868 | 38.84 | 227 | 119 |
|  | TT Zucchini MOCK Ta26 vs TT Zucchini WMV Ta26 | 17169/27868 | 61.60 | 279 | 256 |
|  | TT Zucchini MOCK Ta32 vs TT Zucchini WMV Ta32 | 16332/27868 | 58.60 | 101 | 497 |
|  | TS Zucchini MOCK Ta20 vs TS Zucchini MOCK Ta26 | 16722/16722 | 60 | 1772 | 2741 |
|  | TS Zucchini MOCK Ta32 vs TS Zucchini MOCK Ta26 | 16101/16101 | 57.77 | 212 | 108 |
|  | TT Zucchini MOCK Ta20 vs TT Zucchini MOCK Ta26 | 15864/15864 | 56.92 | 2827 | 3337 |
|  | TT Zucchini MOCK Ta32 vs TT Zucchini MOCK Ta26 | 15061/15061 | 50.04 | 652 | 975 |

### Functional annotation, and GO-Term enrichment analysis

Functional annotation of specific DEGs found in each variety of melon and zucchini under the combination of temperature and virus infection was extracted from AmiGO 2 (https://amigo.geneontology.org/amigo/landing). Using the taxonomy ID 3656 and 3664 for melon and zucchini respectively, the visualization platform of iDEP 2.0 (http://bioinformatics.sdstate.edu/idep/) and ShinyGO 0.77 (http://bioinformatics.sdstate.edu/go/) were used to perform Gene Ontology (GO) enrichment analysis and the protein–protein interaction network, using all available gene sets with an FDR cutoff of 0.05 and log_2_FC <-1 or >1. The top 15 pathways were considered.

### Identification of orthologous genes

To identify orthologous genes between melon and zucchini under different temperature conditions and viral infection, reference genomes were obtained from the public Cucurbit Genomics Database (CuGenDB) (http://cucurbitgenomics.org/) (Yu *et al*., 2023). We selected the well-annotated reference genomes of Melon (DHL92) v3.6.1 and *Cucurbita pepo* subsp. Pepo. The identification of orthologous genes was carried out using the genes involved in the top 15 pathways with the Synteny Viewer tool.

### Statistical analyses

The analysis of the viral load for each plant species was performed with a two-way ANOVA. The viral accumulation data were transformed using a logarithmic function to achieve normality. The model included the plant species (melon and zucchini) and the temperature as two-level fixed effects. All analyses were performed in JMP software. Plot graphs of the viral RNA accumulation for each isolate and plant species were performed using R software. Significant changes in transcript expression as compared to the control (DEGs) were defined as log_2_ FC <-1 or >1 and adjusted P < 0.05 (negative binomial Wald test followed by Benjamini–Hochberg correction) (Ferreira & Zwinderman, 2006).

### Data availability

All RNA sequencing (3’mRNA-seq) data that support the finding of this study have been deposited in ArrayExpress (EMBL-EBI), with the accession codes E-MTAB-14116 for melon and E-MTAB-14118 for zucchini plant species. The raw data, processed data, and metadata with detailed sample annotations and protocols of this 3’mRNA study is provided.

## RESULTS AND DISCUSSION

### WMV load is significantly influenced by melon and zucchini plants and depends on the plant’s thermotolerance level

To determine to which extent temperature affects viral accumulation in melon and zucchini plants, WMV load was estimated using absolute RT-qPCR in commercial melon (Piel de sapo) and zucchini (Black beauty) cultivars grown at different temperatures (Low: 20/14°C, Medium: 26/20°C, and High: 32/26°C). After 30 dpi, WMV had five-fold greater viral accumulation in zucchini than in melon plants (Fig. S1, A-B; F_1,12_= 193.23 *p* < 0.001), with a significant interaction between temperature and plant species (F_2,12_= 78.09 *p* < 0.001). This viral accumulation pattern may have arisen because of host specificity and limited viral fitness of WMV in melon plants at high temperatures. Many studies have shown that viral titers can be influenced by temperature. For example, turnip mosaic virus (TuMV) and plum pox virus (PPV) show lower accumulation at high temperatures, with milder symptoms (Szittya *et al*., 2003; Aguilar *et al*., 2015; Chung *et al*., 2015). In contrast, TuMV has been observed to exhibit more severe symptoms in Chinese cabbage at 28°C, which correlates with higher viral accumulation (Chung *et al*., 2015). Though, plants may experience a slowdown in metabolic processes at lower temperatures, which could affect the efficiency of plant defense mechanisms and, in turn, may affect viral replication and accumulation (Garcia-Ruiz, 2018). Thus, we aimed to evaluate how plant thermotolerance could affect viral accumulation under varying temperatures. Specifically, we examined WMV accumulation in melon and zucchini plants with different levels of thermotolerance (TT: Thermo-tolerant and TS: Thermo-susceptible) under three temperature ranges (L, M, and H). In melon plants, the WMV load decreased significantly in the TS cultivar as the temperature increased, with a 7-fold reduction in the high-temperature range (H) (F_2,6_= 6.12; *p* = 0.003). In contrast, the WMV load in the TT melon cultivar did not vary significantly across temperature ranges (F_2,6_= 0.11; *p* = 0.89). However, in zucchini plants, the WMV load was generally higher than in melon, and its accumulation pattern was reversed. In TS zucchini plants, the WMV load remained relatively at a similar level across the three temperature ranges in the TS plants (F_2,6_= 4.62; *p* = 0.061), whereas in the TT zucchini plants, it significantly increased in the H-temperature range (F_2,6_= 19.90; *p* = 0.002) (Fig. 2 A and B (TS vs TT); two-way ANOVA analysis). These results indicated different responses to heat-stress between TS and TT cultivars in both cucurbit species, whereas WMV load had a relative decrease with increasing temperature in TS plants, while TT plants exhibited certain resilience under temperature changes. These responses suggest that plant susceptibility to temperature affects viral accumulation under heat-stress. In this sense, it has been reported that RNA silencing-mediated defence can be inhibited at low temperatures (Szittya *et al*., 2003; Chellappan *et al*., 2005), which in turn can increase virus accumulation. However, temperature can also affect phytohormone-mediated defense pathways, influencing RNA silencing and virus-encoded silencing suppressors (Lewsey *et al*., 2010). Other studies have shown that high temperatures may increase the susceptibility of tomato plants to tomato yellow leaf curl virus (TYLCV), Arabidopsis to turnip mosaic virus (TuMV), or potato to PVY (Prasch & Sonnewald, 2013; Ghandi *et al*., 2016; Fesenko *et al*., 2021). In the latter case, and in contrast to our results, these studies found that rising temperatures lead to greater accumulation of PVY in thermo-sensitive than in thermo-tolerant plants, possibly due to the reduction of pathogenesis-related proteins (*salicylic acid (SA)-mediated plant defense*) (Makarova *et al*., 2018; Spechenkova *et al*., 2021). It is likely that an increase in temperature can reduce plant resistance, as the case for tobacco mosaic virus (TMV) and potato potexvirus X (PVX), where an increase in temperature led to a reduction in the induced hypersensitivity response (Wang *et al*., 2009). However, it has been also reported that temperatures above 28°C can induce a suppression of virus-induced HR-type necrosis caused by TMV (Király *et al*., 2008). In addition, increased temperature and faster symptom development could also be correlated with virus accumulation, as observed for peanut stunt virus in *N. benthamiana* (Obrępalska-Stęplowska *et al*., 2015), PVY in potato (Makarova *et al*., 2018), and capsicum chlorosis virus in pepper (Tsai *et al*., 2022a). Taken together, our results suggest that temperature can influence WMV accumulation in melon and zucchini plants, based on their thermotolerance level. This outcome could be related to differences in the expression of genes and proteins involved in stress responses, which could indirectly affect viral replication, movement, and/or accumulation within the plant.

**Figure 2.**
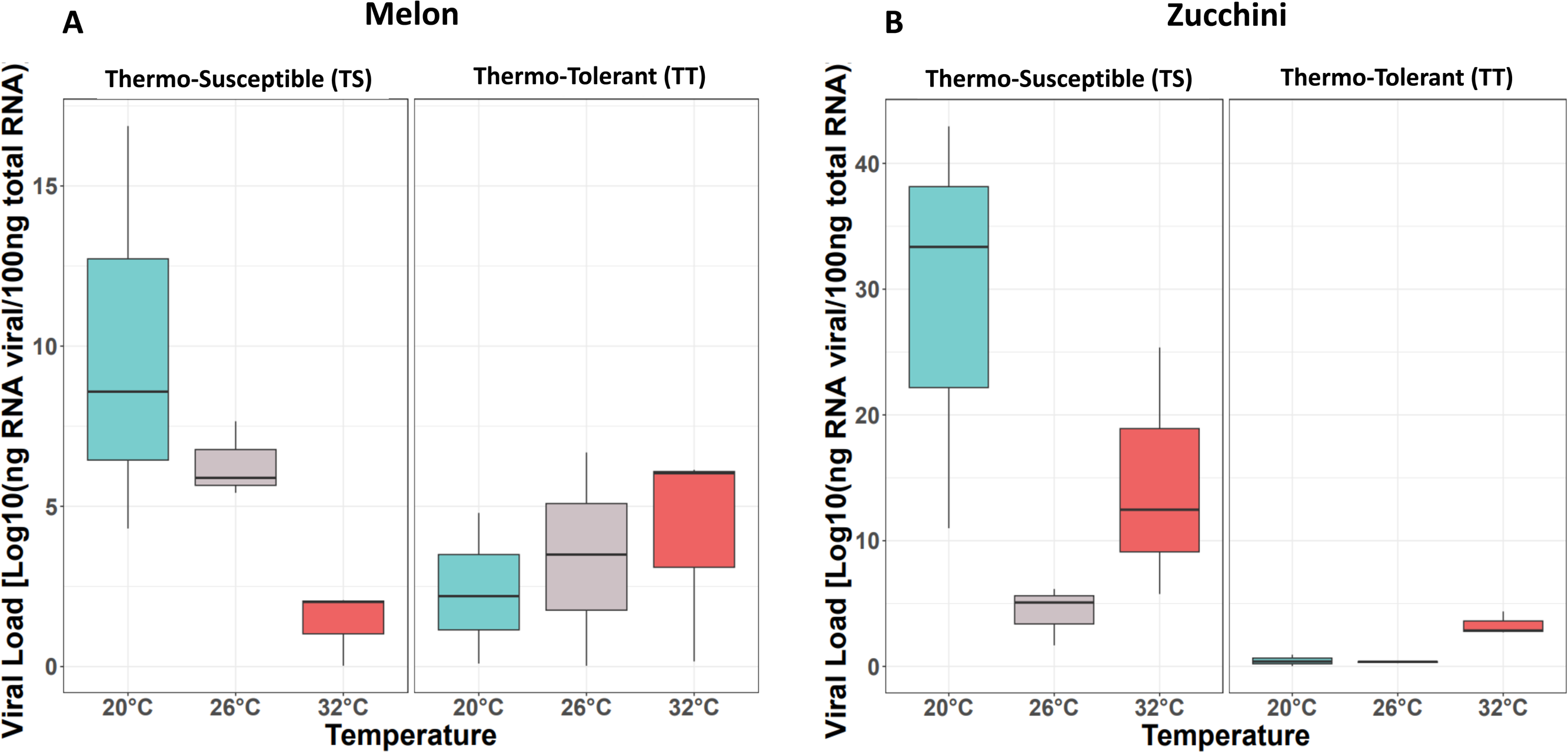
Viral load (mean and SE error bars, n = 3) of WMV infectious clones (MeWMV7) in thermo-susceptible (TS) and -tolerant (TT) melon and zucchini plants at 30 dpi under three different growth temperatures: 20°C/16°C (Low, blue color), 26°C/20°C (Medium, grey color) or 32°C/24°C (High, red color). Viral RNA accumulation was inferred by quantitative RTq-PCR. RNA transcripts of the P1 gene were used in serial dilutions (10-fold) to generate external standard curves. The RNA concentration in each sample (ng of viral RNA per 100 ng of total RNA) was estimated from the cycle threshold (Ct) values obtained from each independent biological assay, with three biological replicates at each time point.

### Significant differences in the number and distribution of DEGs across temperature ranges and WMV infection between melon and zucchini plants

To examine the effect of WMV infection and temperature variations (L, M, and H) on the gene response of melon and zucchini, with temperature tolerance (TT and TS), we carried out a differential 3’mRNA-seq approach of plants under each temperature condition in the presence or absence of WMV. The reads obtained from each replicate sample were normalized and analyzed using principal component analysis (PCA). Overall, the PCA clustered mock samples separately from virus-infected samples, as well as by temperature conditions within the TS and TT melon and zucchini samples (Fig. S2: A-B). After the bioinformatic processing of the total differentially expressed genes (DEGs), and the subsequent analysis using the criteria *p* < 0.05 and log_2_ FC ≤ −1 or ≥ 1, we found a total of 14,955 differentially expressed genes (DEGs) at low (337 overexpressed and 119 underexpressed), 14,882 at medium (115 overexpressed and 220 underexpressed), and 15,351 at high (519 overexpressed and 1,102 underexpressed) temperatures in TT melon while 15,150 differentially expressed genes (DEGs) at low (118 overexpressed and 110 underexpressed), 10,643 at medium (50 overexpressed and 115 underexpressed), and 10,544 at high (133 overexpressed and 280 underexpressed) temperatures in TS melon, out of 37,254 annotated genes in melon. In zucchini plants, we found a total of 21,938 DEGs (109 overexpressed and 299 underexpressed) at low, 15,963 at medium (183 overexpressed and 295 underexpressed), 16,281 at high temperatures (366 overexpressed and 640 underexpressed) in TS zucchini, and 10,825 DEGs (227 overexpressed and 119 underexpressed) at low, 17,169 at medium (279 overexpressed and 256 underexpressed), 16,332 at high temperatures (101 overexpressed and 497 underexpressed) in TT zucchini, out of 27,868 annotated genes in zucchini. For more information on the total number of genes overexpressed and underexpressed for each condition and crop, refer to Supplementary Tables 1 - 3. Thus, zucchini plants had approximately 2.6-fold higher percentage of DEGs in melon (FRatio= 17.9602 and Prob>F= 0,0028). In TS and TT plants, the percentage of DEGs in melon ranged from 10,544 (28%) to 15,351 (41%), respectively out of 37,254 annotated genes, whereas in zucchini, it ranged from 10,825 (39%) to 21,938 (79%), respectively out of 27,868 annotated genes (Table 1). Furthermore, the % DEGs were similar in TT melon and zucchini plants across temperatures (40% and 53%, respectively). In TS melon and zucchini plants, the highest percentage of DEGs occurred at lower temperatures in TS melon (41%) and TS zucchini (79%) plants. Note that this high % of DEGs in TS at low temperatures may be related to the greater accumulation of WMV, suggesting specific transcriptional responses of TS to the combination of temperature and virus infection. This is consistent with previous studies that have shown that low-temperature conditions make plants more susceptible to viruses because of the inhibition of RNA silencing-mediated defence by controlling siRNA generation (Szittya *et al*., 2003; Tenllado & Canto, 2020).

We then sought to identify unique genes under both viral infection and temperature-stress conditions. In both, the TS and TT of melon and zucchini samples, the highest number of unique DEGs was observed under the condition of WMV infection and high-temperature, whereas only a few (2-4) DEGs (Table S1) were found to be common for all conditions (Fig. 3: A-D). Specifically, in TS and TT melon plants, 205 and 711 genes were unique under WMV+H conditions, respectively; 21 and 40 genes were unique at WMV+M, respectively; and 83 and 88 genes at WMV+L, respectively. In TS and TT zucchini plants, 306 and 121 genes were unique at WMV+H, respectively; 126 and 27 genes at WMV+M, respectively; and 153 and 61 genes were unique at WMV+L, respectively (for a more detailed information, see Fig. S3). This indicated that the effect of WMV infection at the high temperature range can be associated with a greater number of unique DEGs in TS and TT from both plant species when compared to the medium and low temperature ranges. It has been suggested that some plant virus infections can mitigate the detrimental effects of abiotic stress in plants (Gorovits *et al*., 2019; Aguilar & Lozano-Duran, 2022; Mishra *et al*., 2022). For example, TYLCV infection in tomato plants can suppress the heat shock response, alleviate the response of plant cells to heat stress, prevent cell death, and allow adaptation of the plant to high temperatures and water deficit (Corrales-Gutierrez *et al*., 2020; Gorovits *et al*., 2022). Alternatively, TuMV infection has been shown to reduce stomatal conductance, leading to altered expression levels of ABA homeostasis genes including biosynthesis and catabolism, eventually improving drought tolerance in Arabidopsis plants (Manacorda *et al*., 2021). It is therefore that specific signaling pathways can be involved in these biotic and abiotic processes, and further research is needed to identify potential targets and fully understand this complex interplay.

**Figure 3.**
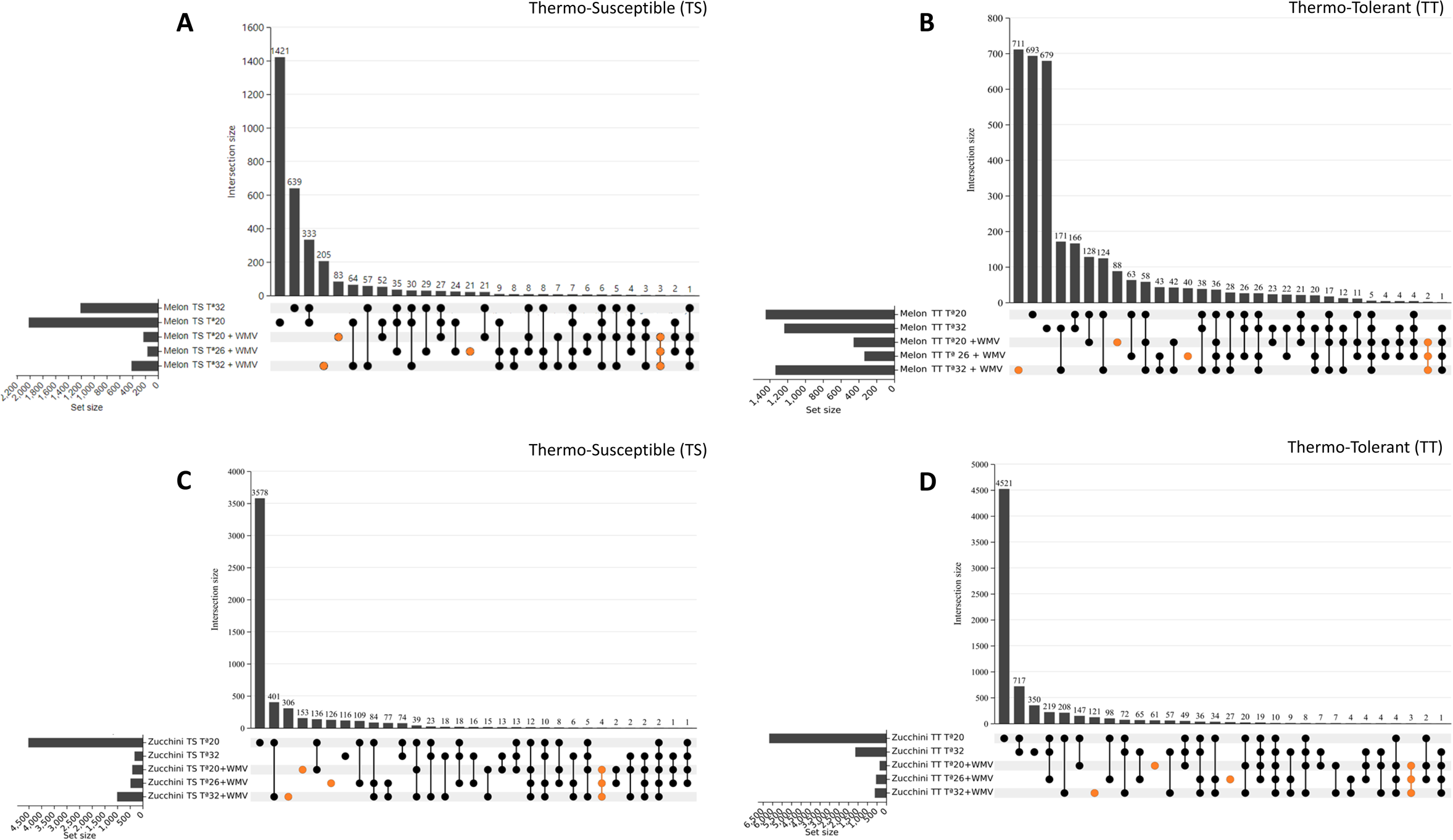
Upset plot displaying the intersections between the sets of DEGs found under single or combined stresses in thermo-susceptible melon **(A)**, thermo-tolerant melon **(B)**, thermo-susceptible zucchini **(C)** and thermo-tolerant zucchini **(D)**. Upregulated and downregulated genes for each variety, crop and condition were put in the same set for the specific DEGs subtraction under the combination of temperature and WMV infection. The bar plot at the top represents the overlap of DEGs under each condition. The horizontal bar on the left represents the number of significant DEGs in each condition. The black circles and lines at the bottom show the categories for which the overlap was calculated, as indicated in the bar chart on the top. The orange circles highlight the DEGs exclusive to each condition in combination with both stresses (WMV+L, WMV+M and WMV+H) and the overlap of significant DEGs observed in the three conditions.

### Changes in biological processes and molecular functions in response to WMV infection and temperature

We next examined the genes that likely contributed to this differentiation of DEGs, which could be potentially associated with both virus infection and heat-stress. Since functional genomic studies in cucurbits are ongoing and there are still transcripts that are poorly annotated in the KEGG database, we used Gene ontology (GO) enrichment to assign DEGs with biological processes and molecular functions. GO analysis was performed for both up- and down-regulated sets of unique genes, which were selected according to the criteria *p* < 0.05 and log_2_FC ≤ −1 or ≥ 1, including an FDR adjustment of 0.05, as the threshold of significance. Among all of them (Table S2-5), the top 15 pathways (up- and down-regulated) for each temperature condition were ranked by *p*-value (Fig. 4, melon) and (Fig. 5, zucchini), with more detail information into the Tables 4 and 5. We found that a considerable % of the DEGs were related to biotic and/or abiotic stress in TS and TT melon and zucchini samples. In particular, in TT samples, 32%, 23%, and 48% of the DEGs were found at low, medium, and high temperatures, respectively. Similarly, in TS samples, 41%, 50%, and 41% were found at low, medium, and high temperatures, respectively. This suggested that temperature had a greater impact on the DEGs in TS samples. Note that the remaining enriched GO-Terms were related to other processes that are not currently associated with biotic and abiotic stress. Among the GO terms related to biotic stress, some annotated common DEGs were *defence response* (GO:0006952), *Hsp70 protein binding* (GO:0030544), *plant-pathogen interaction* (Path:cmo04626), and those related to *jasmonic acid metabolism* (GO:0009694 and GO:0009695) (Table S6-7). These particular genes have been reported to be involved in biotic stress responses (Jones & Dangl, 2006; Mittler *et al*., 2012; Wasternack & Strnad, 2018). Similarly, among the GO terms related to temperature stress, the annotated DEGs were *responses to radiation or light stimulus* (GO:0009314 and GO:0009416) or *cellular copper ion homeostasis* (GO:0006878) and were reported to be involved in abiotic stress (Wang, 2005; Yruela, 2005; Hideg *et al*., 2013). Furthermore, there were also GO terms related to both abiotic and biotic stressors, such as *protein folding, secretion, processing* (GO: 0006457, GO:0009306, and GO:0016485), and *lipid metabolism* (GO:0006631) (Queitsch *et al*., 2000; Narayanan *et al*., 2016). The analysis of TT melon plants revealed that *ATPase-coupled intramembrane lipid transporter activity* (GO:0140326 and GO:0140303) and the *glyoxylate cycle* (GO:0006097) were enriched with upregulated and downregulated genes, respectively, at low-temperature range (Fig. 4A and 4D). Specifically, in the case of ATPase-coupled intramembrane lipid transporter activity, GO-Term was strongly related to other highly significant factors, such as *lipid transport* (GO:0140303 and GO:0005319) (Fig. 5A). At medium-temperature range, *isocitrate metabolic process* (GO:0006102) and *phagosome* (Path:cmo04145) were enriched with upregulated and downregulated genes, respectively (Fig. 4B and 4E) that were associated with and involved in similar metabolic pathways (Fig. 5B). While a large number of GO Terms were enriched at high-temperature range, highlighting the *glycine catabolic process* (GO:0006546) and *serine family amino acid catabolic process* (GO:0009071) with upregulated genes (Fig. 4C) that were associated with and involved in similar metabolic pathways (Fig. 5C), as well as *jasmonic acid metabolic process* (GO:0009694) with downregulated genes (Fig. 4F) and related to *jasmonic acid biosynthesis* (GO:0009695) (Fig. 5C). On the other hand, in TS melon, the *carotene metabolic process* (GO:0016120) and *Golgi to plasma membrane transport* (GO:0006893) were found to be upregulated and downregulated genes, respectively, at low-temperature range (Fig. 4G and 4J). *The carotene metabolic process* has been highly related to other GO terms such as *carotene biosynthetic process,* and some of which are associated with *cellular alcohol* (GO:0044107) and *ergosterol process* (GO:0008204 and GO:0006696) (Fig. 5D). At medium-temperature range, *protein import into peroxisome docking* (GO:0016560) and *phenylalanine metabolism* (Path:cmo00360) were enriched with upregulated and downregulated genes, respectively (Fig. 4H and 4K) that were associated with and involved in similar metabolic pathways (Fig. 5E).

**Figure 4.**
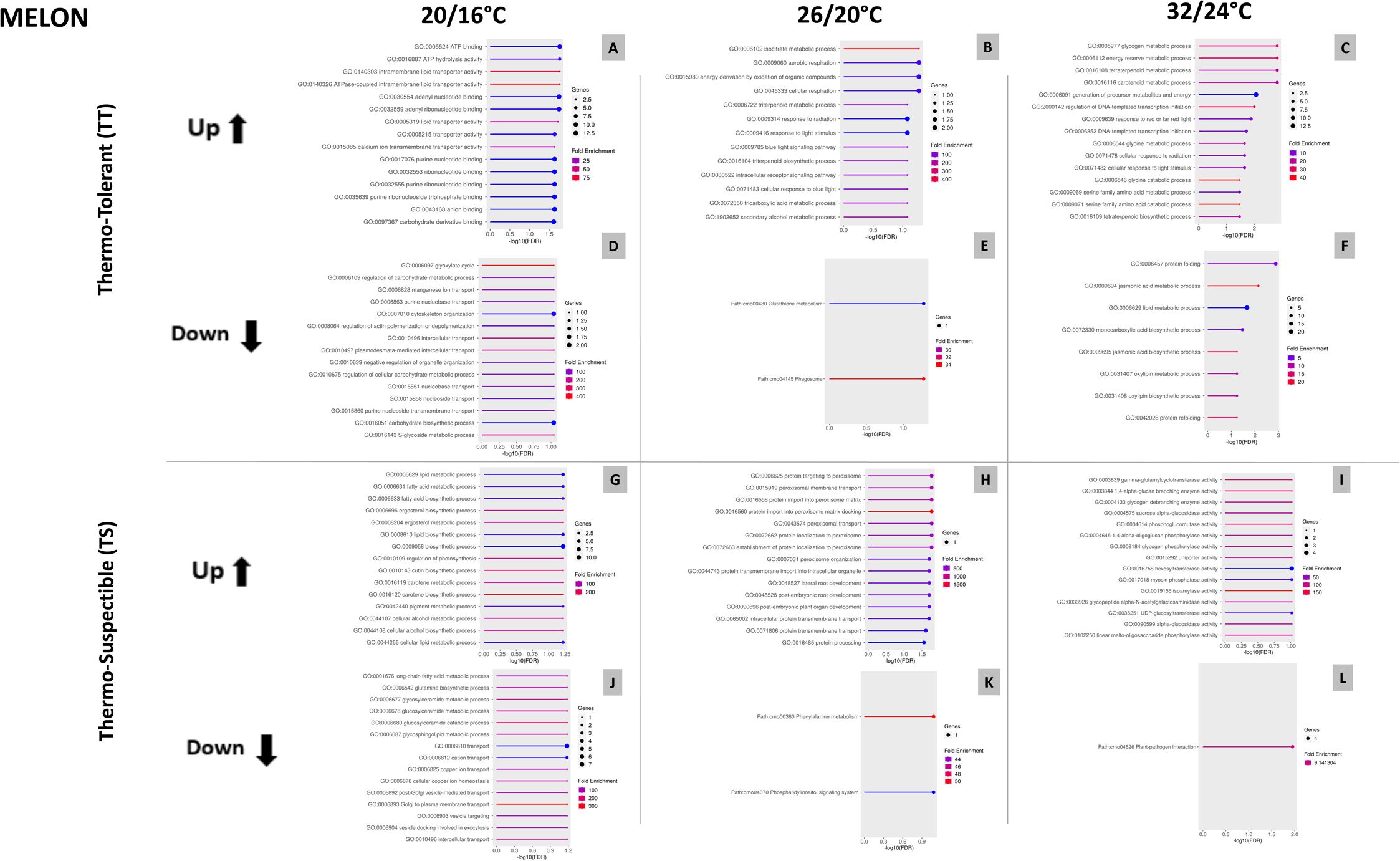
The Gene Ontology (GO) enrichment analysis for the top of 15 GO enrichment of specific DEGs (using the criteria *p* < 0.05 and Log_2_FC ≤ −1 or ≥ 1) in melon plant varieties under the combination of Temperature+WMV infection based on Biological Process and Molecular Function. GO enrichment profiles are shown for thermo-tolerant upregulated genes at low **(A)**, medium **(B)** and high **(C)** temperatures and downregulated genes at low **(D)**, medium **(E)** and high **(F)** temperatures. GO enrichment profile are shown for thermo-susceptible upregulated genes at low **(G)**, medium **(H)** and high **(I)** temperatures and downregulated genes at low **(J)**, medium **(K)** and high **(L)** temperatures.

**Figure 5.**
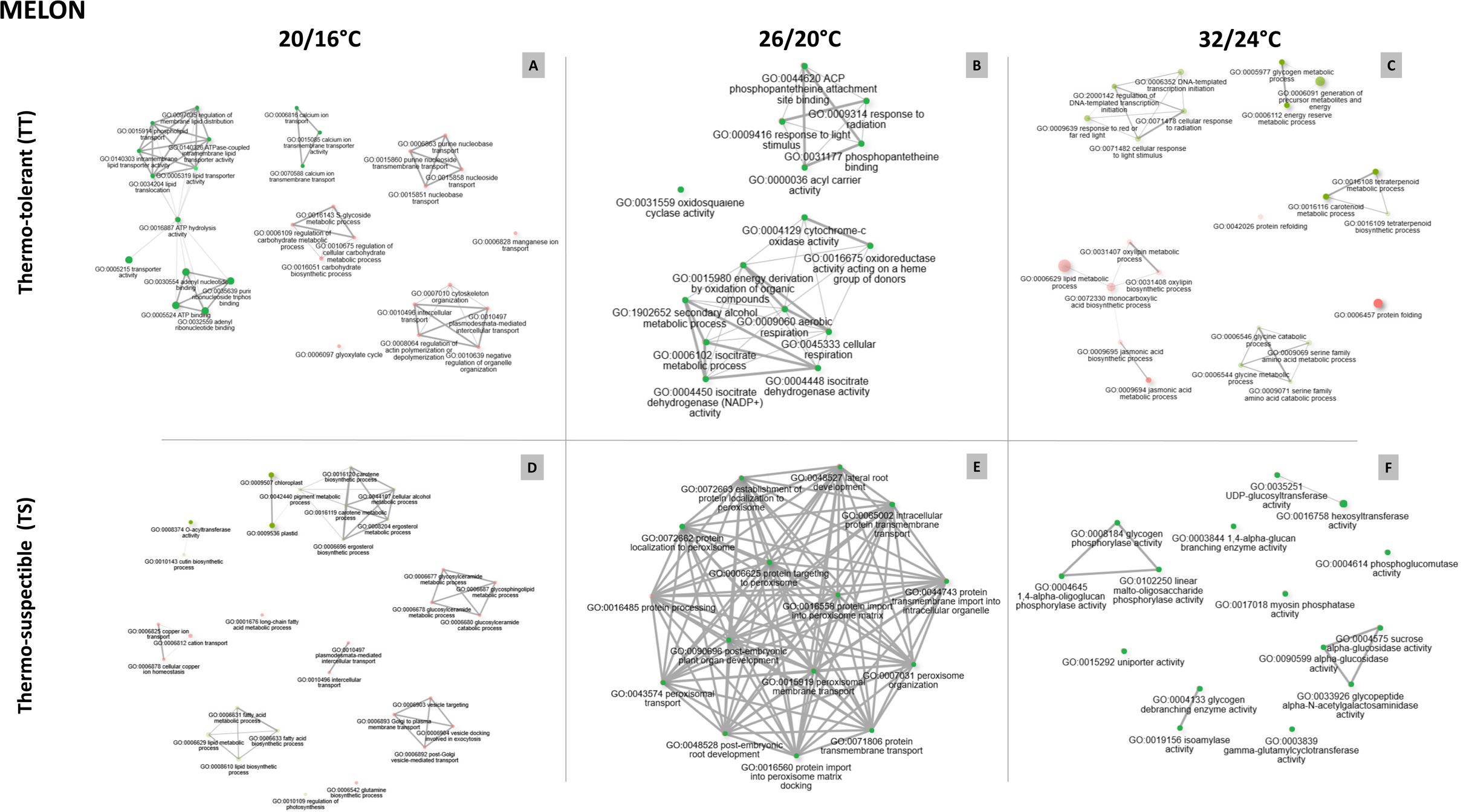
Predicted functional GO pathway interaction networks based on Molecular Function and Biological Process of specific DEGs under the combination of Temperature+WMV infection in melon plant varieties (FDR cutoff: 0.3). The size of the circles is proportional to the number of genes related to that pathway and the color intensity is related to the pathway P-value significance level (P < 0.05). The green and red colors represent upregulated and downregulated genes, respectively. Networks are shown for thermo-tolerant (TT) at low **(A)**, medium **(B)** and high **(C)** temperatures and thermo-susceptible (TS) at low **(D)**, medium **(E)** and high **(F)** temperatures. The raw data used for this figure are listed in the Table S6.

While, at high-temperature range, *isoamylase activity* (GO:0019156), which has been associated with *glycogen debranching enzyme activity* (GO:0004133) (Fig. 5F), and *plant-pathogen interaction* (Path:cmo04626) was found to be upregulated and downregulated genes, respectively (Fig. 4I and 4L).

The analysis of the TT zucchini plants showed that *calmodulin-lysine N-methyltransferase activity* (GO:0018025) and *fluoride transmembrane transporter activity* (GO:1903425) were enriched with upregulated genes (Fig. 6A), and the *farnesol dehydrogenase activity* (GO:0047886), *nicotinate N-methytransferase activity* (GO:0008938), and *biotin carboxylase activity* (GO:0004075) were enriched with downregulated genes at low-temperature range (Fig. 6D), none of which was associated with any of the top 15 pathways (Fig. 7A). At medium-temperature range, *COPI coating of Golgi vesicle* (GO:0048205) a*nd iron assimilation by chelation and transport* (GO:0033214) were enriched with upregulated and downregulated genes, respectively (Fig. 6B and 6E) that were associated with and involved in similar metabolic pathways (Fig. 7B). While at high-temperature range, the *nucleoside transmembrane transport* (GO:1901642), which has been associated with *nucleoside transmembrane transporter activity* (GO:0005337) (Fig. 7C), and the *triglyceride metabolic process* (GO:0006641), which has been associated with *glycerol-phosphate biosynthesis process* (GO:0046167) (Fig. 7C), were both enriched with upregulated and downregulated genes, respectively (Fig. 6C and 6F). In TS zucchini plants, the analysis showed that the *positive regulation of cellular response to phosphate starvation* (GO:0080040) and *trehalose metabolic process* (GO:0005991), which have been associated with *alpha, alpha-trehalase activity* (GO:0004555) (Fig. 7D), were both enriched with upregulated and downregulated genes, respectively, at low-temperature range (Fig. 6G and 56J). At medium-temperature range, *mitocondrial respiratory chain complex III assembly* (GO:0034551); and response to *light intesity* (GO:0009642), *copper ion* (GO:0046688) and *mitocondrial ribosomal subunit assembly* (GO:1902775) were enriched with upregulated and downregulated genes, respectively (Fig. 6H and 6K) that were associated with and involved in similar metabolic pathways (Fig. 7E). While at high-temperature range, the *fructose and glucose transport* (GO:0015755, GO:0005353 and GO:1904659), which have been strongly associated with similar metabolic pathways (Fig. 7F), and the *protein N-linked glycosylation via asparagine* (GO:0018279), were also found to be upregulated and downregulated genes, respectively (Fig. 6I and 6L). The identification of these genes suggests that certain biological processes may be linked to both abiotic and biotic stresses. Although it should be noted that our results could be influenced by the specific temperature range and pathogen used in our experimental design, it is evident that the combination of these stress factors can play an important role in plant physiology. It is also worth mentioning that despite the cucurbit genomics database (CuGenDB) is a pivotal and valuable resource for advancing research for comparative and functional genomic studies (Zheng *et al*., 2018), functional genomic studies in non-model crops are usually challenging, and further research is needed to elucidate the specific genes underlying abiotic and biotic stresses.

**Figure 6.**
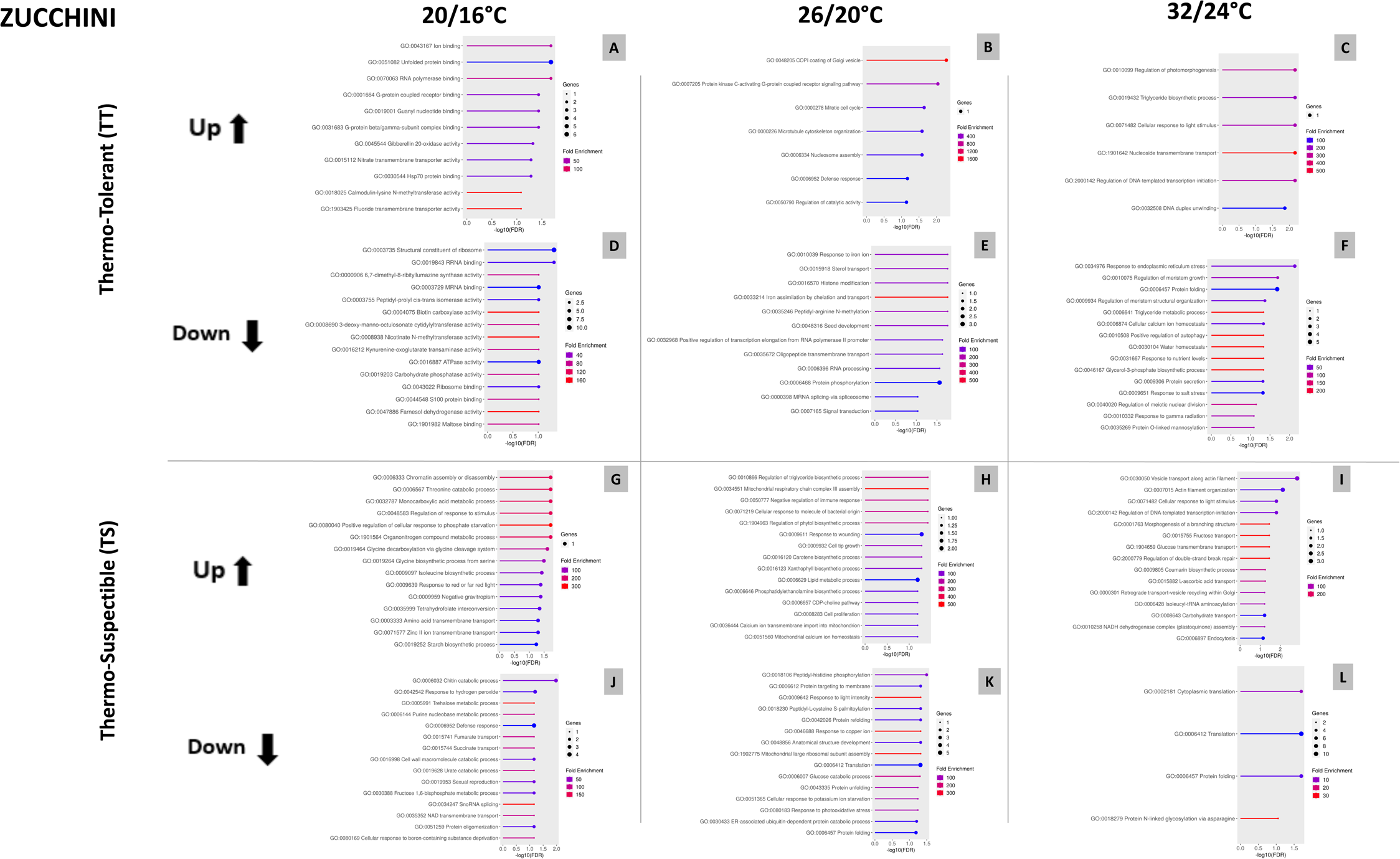
The Gene Ontology (GO) enrichment analysis for the top of 15 GO enrichment of specific DEGs (using the criteria *p* < 0.05 and Log_2_FC ≤ −1 or ≥ 1) in zucchini plants varieties under the combination of Temperature+WMV infection based on Biological Process and Molecular Function. GO enrichment profiles are shown for thermo-tolerant upregulated genes at low **(A)**, medium **(B)** and high **(C)** temperatures and downregulated genes at low **(D)**, medium **(E)** and high **(F)**. GO enrichment profiles are shown for thermo-susceptible upregulated genes at low **(G)**, medium **(H)** and high **(I)** temperatures and downregulated genes at low **(J)**, medium **(K)** and high **(L)** temperatures.

**Figure 7.**
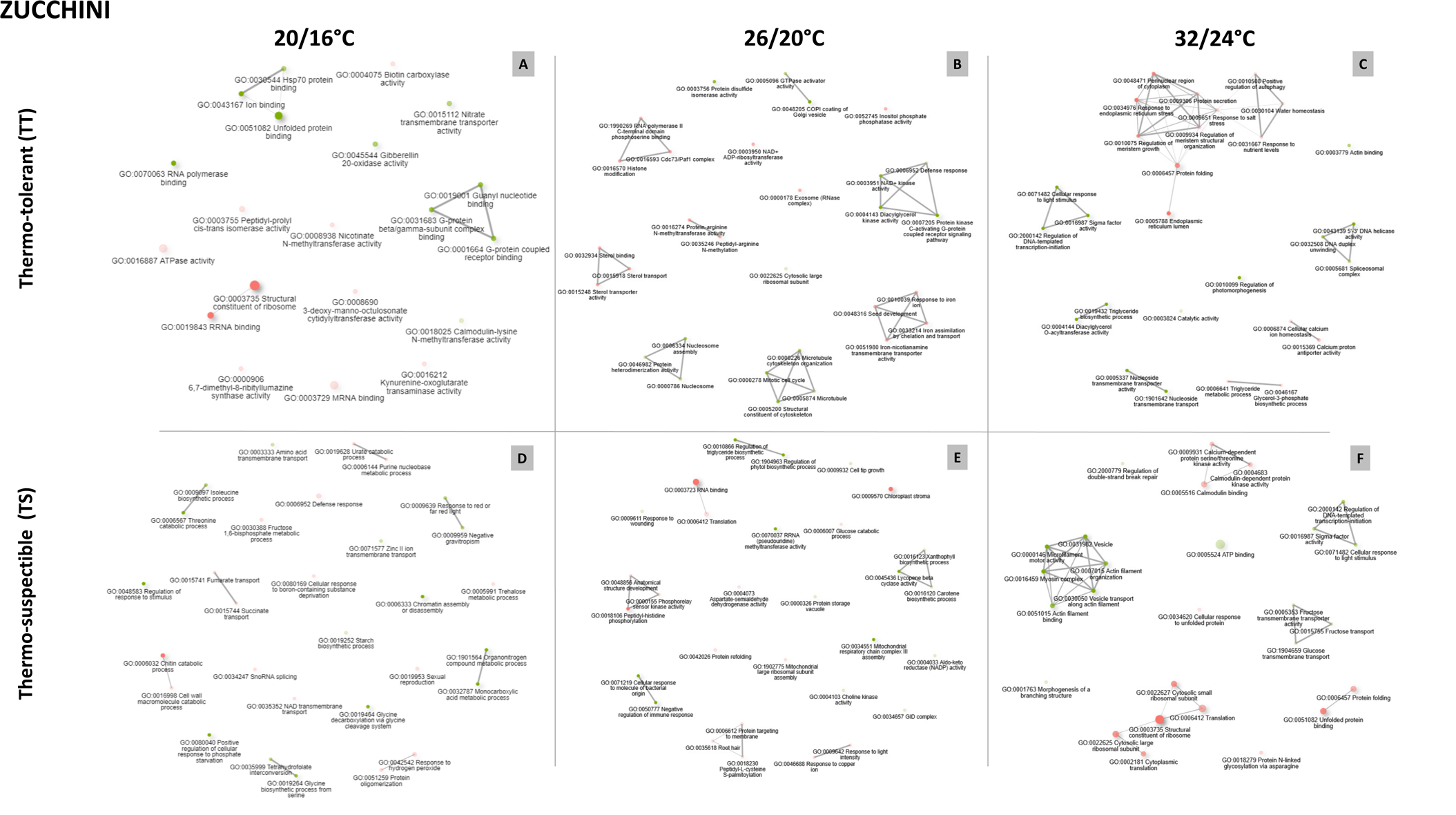
Predicted functional GO pathway interaction networks based on Molecular Function and Biological Process of specific DEGs under the combination of Temperature+WMV infection in zucchini plants varieties (FDR cutoff: 0.3). The size of the circles is proportional to the number of genes related to that pathway and the color intensity is related to the pathway P-value significance level (P < 0.05). Green and red colors represent upregulated and downregulated genes, respectively. Networks are shown for thermo-tolerant (TT) at low **(A)**, medium **(B)** and high **(C)** temperatures and thermo-susceptible (TS) at low **(D)**, medium **(E)** and high **(F)** temperatures. The raw data used for this figure are listed in the Table S7.

### Identification of unique orthologous genes in thermo-susceptible and thermo-tolerant melon and zucchini plants

A comprehensive overview and identification of the orthologous genes across these melon and zucchini plant species can indeed provide critical insights into the genetic basis of stress response pathways and could point to specific pathways involved in the combination of temperature and viral infection stress. To this end, orthologous genes were identified among the top 15 pathways for each temperature in the TS and TT melon and zucchini plants. At high-temperature range, one unique orthologous gene was found between TT melon and zucchini plants. In melon, this gene is referenced as MELO3C023308.2 (log_2_(FoldChange)=1.709) and the orthologous gene in zucchini is refered to Cp4.1LG05g12560 (log_2_(FoldChange)=1.036), both being overexpressed and exclusive owing to the condition of high-temperature tolerance. This melon gene is annotated with the following GO-Terms: GO:0009658, GO:0048512, GO:0045893, GO:0080167, GO:0010099 and GO:0009639, which are related to chloroplast organization, circadian behavior, positive regulation of transcription, response to karrikin, regulation of photomorphogenesis under the biological process category and response to red or far red light, respectively. In the protein–protein interaction network, it was directly related to regulation of *DNA-templated transcription initiation* (GO:2000142), *cellular response to radiation* (GO:0071478) and *light stimulus* (GO:0071482) (Fig. 5C). In zucchini, the gene mentioned is annotated under GO:0010099 term, which is related to the regulation of photomorphogenesis under the biological process category. This is consistent with studies on *Arabidopsis thaliana,* which showed that thermomorphogenetic effects occur in response to high temperatures (Nomoto *et al*., 2012; Yamashino *et al*., 2013). These thermomorphogenetic responses are typical of thermo-tolerant varieties, making them attractive for crop breeders aiming to develop plants that can withstand climate warming (Quint *et al*., 2016). Additionally, these genes encode F-Box proteins, known to play pivotal roles in regulating the expression of genes involved in plant defense response by recognizing and reacting to pathogen-associated molecular patterns (PAMPs) and effector-triggered immunity (ETI). They are involved in the regulation of protein stability during immune responses, ensuring efficient defense against pathogens, as well as in the degradation of misfolded or damaged proteins accumulated during temperature stress, thus maintaining cellular homeostasis and regulatory proteins that modulate heat stress and hypersensitive resistance responses (Xiao & Jang, 2000). At medium-temperature range, no orthologous genes were found in TT or TS between plant species. And at low-temperature range, one unique orthologous gene was found among the TS melon and zucchini plants. The melon gene is referenced as MELO3C024920.2 (log_2_(FoldChange)=-1.010) and the orthologous gene in zucchini was found with reference Cp4.1LG06g08450 (log_2_(FoldChange)=1.258), both being exclusive due to the condition of low-temperature tolerance and being underexpressed and overexpressed, respectively. This melon gene is annotated with the following GO-Terms: GO:0030001, GO:0055085, GO:0055114, GO:0071577, GO:0006810, and GO:0006812, which are related to metal ion transport, transmembrane transport, oxidation-reduction process, and zinc II ion transmembrane transport under the biological process category, transport and cation transport, respectively. In the protein–protein interaction network, it was directly related to *cellular copper ion homeostasis* (GO:0006878) and *transport* (GO:0006825) (Fig. 5D). In zucchini, the gene mentioned is annotated under the GO:0030001, GO:0055085, GO:0055114 and GO:0071577 which are related to metal ion transport, transmembrane transport, oxidation-reduction process under the biological process category. These genes encode Zinc transporters, which have been shown to be involved in the regulation of various signaling pathways that are crucial for plant growth and development, making it a key player in plant stress responses. Zinc transporters regulate zinc uptake, transport, and distribution, influencing the ability to tolerate and adapt to adverse environmental conditions such as drought, salinity, and heavy metal stress (Ullah *et al*., 2019). plant’s immune response against pathogens by regulating zinc accumulation (Cabot *et al*., 2019). Our results suggest that this orthologous gene may be crucial for maintaining the balance between stress and growth in plants, and further research should be conducted to explore its specific functions in different plant species under combined stresses. Curiously, at temperatures where the accumulation of WMV was lower, no orthologous DEGs were found among zucchini and melon samples. This would suggest that the particular group of DEGs under these conditions can play a significant role in the observed response to the interaction between cucurbit plants and viruses. Further investigation of these potential candidates could lead to the development of cucurbit cultivars with tolerance to both stresses. In conclusion, our study shows that WMV infection elicits an active defense response at the transcriptomic level in both thermo-sensitive and thermo-tolerant cucurbit crops, with the activation of particular signaling pathways that underlie the combined effects of biotic and abiotic stressors.

## ACKNOWLEDGEMENTS

We thank Rosa Rivero (CEBAS-CSIC) for the useful discussions and comments. This work was part of the research project, PID2022-141108OB-I00 funded by MCIN/AEI/10.13039/501100011033/FEDER (EU). CDMR was supported by Fundación Séneca within the PhD programme (SENECA 21417/FPI/20). We acknowledge the support of the publication fee by the CSIC Open Access Publication Support Initiative through its Unit of Information Resources for Research (URICI).

Conflict of interest: The authors declare that they have no conflict of interest.

## Supplementary material

**Figure S1.**
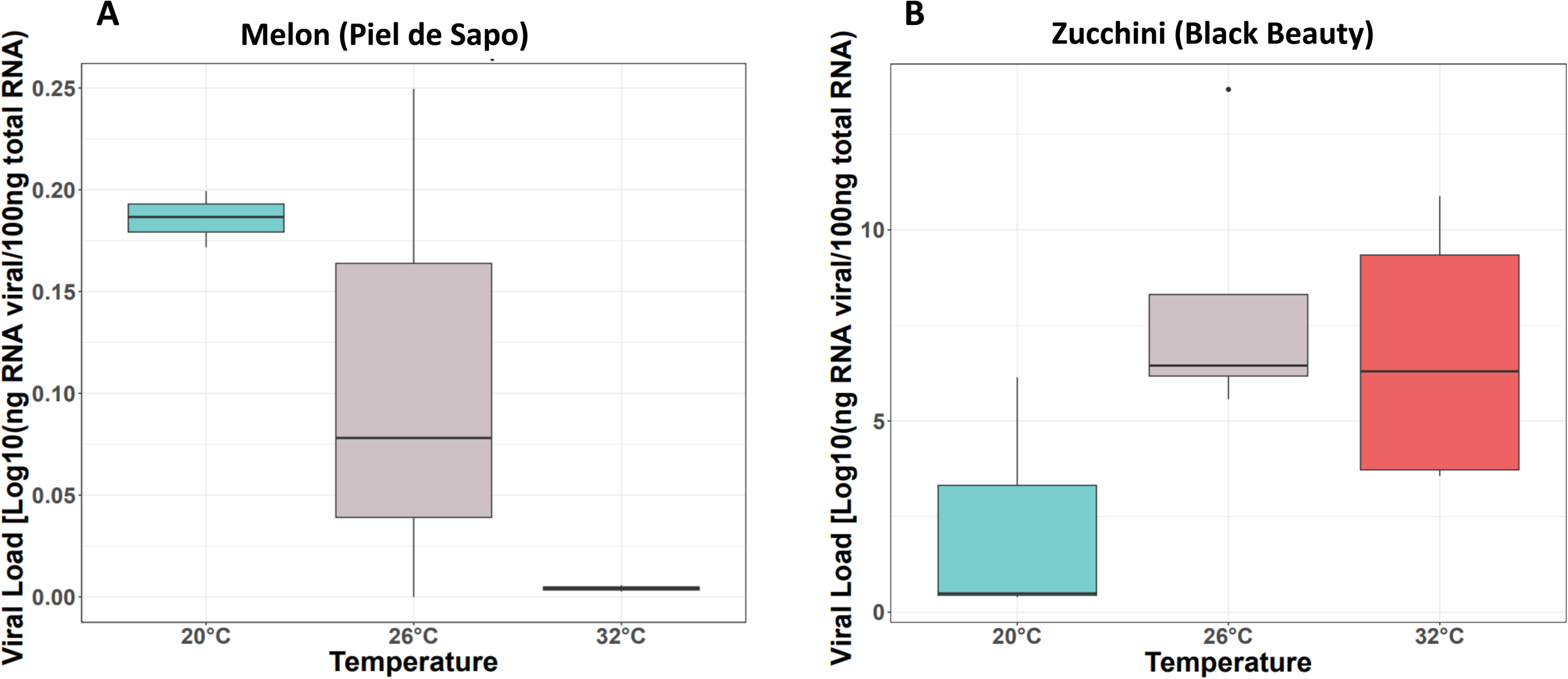
Viral load (mean and SE error bars, n = 3) of WMV infectious clones (MeWMV7) in commercial melon (Piel de sapo) **(A)** and zucchini (Black beauty) **(B)** plants at 30 dpi under three different growth temperatures: 20°C/16°C (Low, blue color), 26°C/20°C (Medium, grey color) or 32°C/24°C (High, red color). Viral RNA accumulation was inferred by absolute quantitative RT-q-PCR. RNA transcripts of the P1 gene were used in serial dilutions (10-fold) to generate external standard curves. RNA concentration in each sample (ng of viral RNA per 100 ng of total RNA) was estimated from the cycle threshold (Ct) values obtained from each independent biological assay, with three biological replicates at each time point.

**Figure S2.**
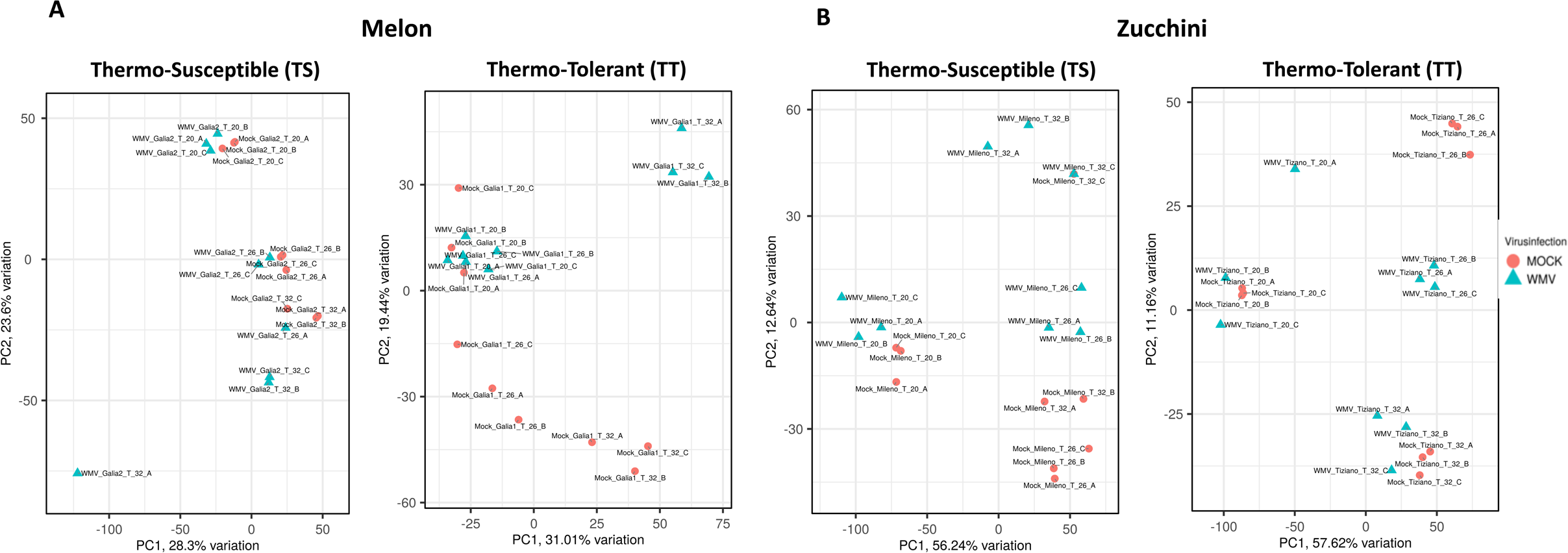
Principal component analysis (PCA) of thermo-susceptible (TS) and -tolerant (TT) melon **(A)** and zucchini **(B)** plants. Mock and infected samples are labeled with the corresponding temperature conditions (20, 26 and 32 °C), including replicates (A, B, and C) and represented by a red circle and a blue triangle, respectively. PCA was performed based on read count data using iDEP 2.01.

**Figure S3.**
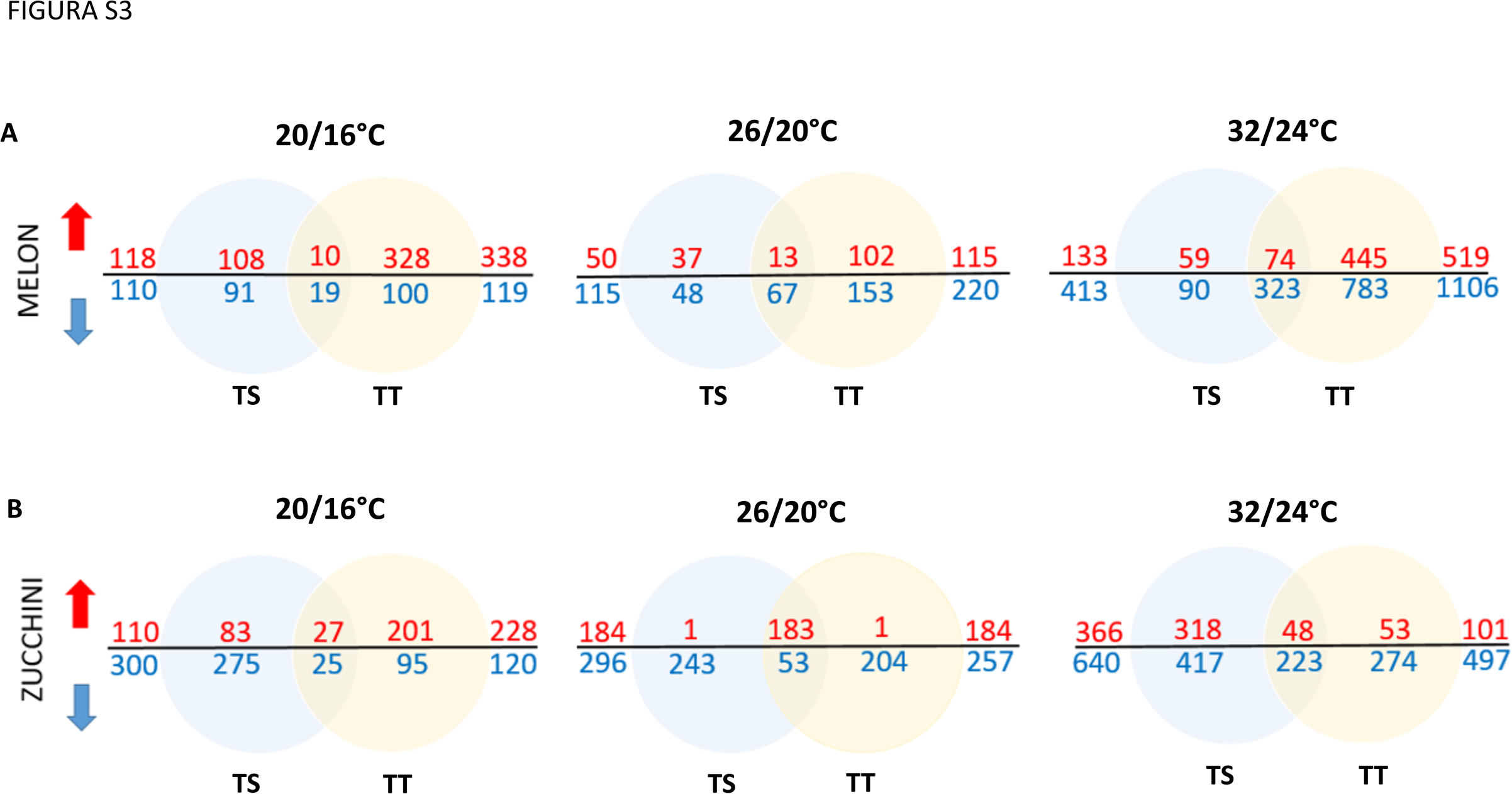
Venn diagrams of the overlap of upregulated and downregulated genes between thermo-tolerant (TT) and thermo-susceptible (TS) **(A)** melon plants and thermo-tolerant (TT) and thermo-susceptible (TS) **(B)** zucchini plants in each stress treatment. Venn diagrams were performed based on the genes list of tables S2 and S3 in R studio.

**Table S1.** Common DEGs observed in the three temperature conditions highlighted in Figure 3 for melon and zucchini varieties are classified by Molecular Function, Biological Process and Cellular Component using the CuGenDB and UniProtKB databases.

**Table S2.** Differentially expressed genes (DEGs) from melon plant varieties under the combination of virus infection and temperatures (p-value < 0.05 and Log_2_FC < −1 o > 1 criteria). Samples were normalized against the Mock-control condition.

**Table S3.** Differentially expressed genes (DEGs) from zucchini plant varieties under the combination of virus infection and temperatures (p-value < 0.05 and Log_2_FC < −1 o > 1 criteria). Samples were normalized against the mock-control condition.

**Table S4.** Differentially expressed genes (DEGs) from melon plants varieties under different temperature conditions (p-value < 0.05 and Log_2_FC < −1 o > 1 criteria). Samples were normalized against the medium (26/20°C) temperature condition.

**Table S5.** Differentially expressed genes (DEGs) from zucchini plants varieties under different temperatures conditions (p-value < 0.05 and Log_2_FC < −1 o > 1 criteria). Samples were normalized against the medium (26/20°C) temperature condition.

**Table S6.** GO enrichment analysis in melon plant varieties, which were specifically regulated under the combination of virus infection and temperatures as shown in Figure 4.

**Table S7.** GO enrichment analysis in zucchini plant varieties, which were specifically regulated under the combination of virus infection and temperatures as shown in Figure 6.

