## Supplementary figures and images for "The combined effect of viral infection and temperature on the gene response of melon and zucchini plants with different levels of temperature tolerance"

### Supp Figure 1

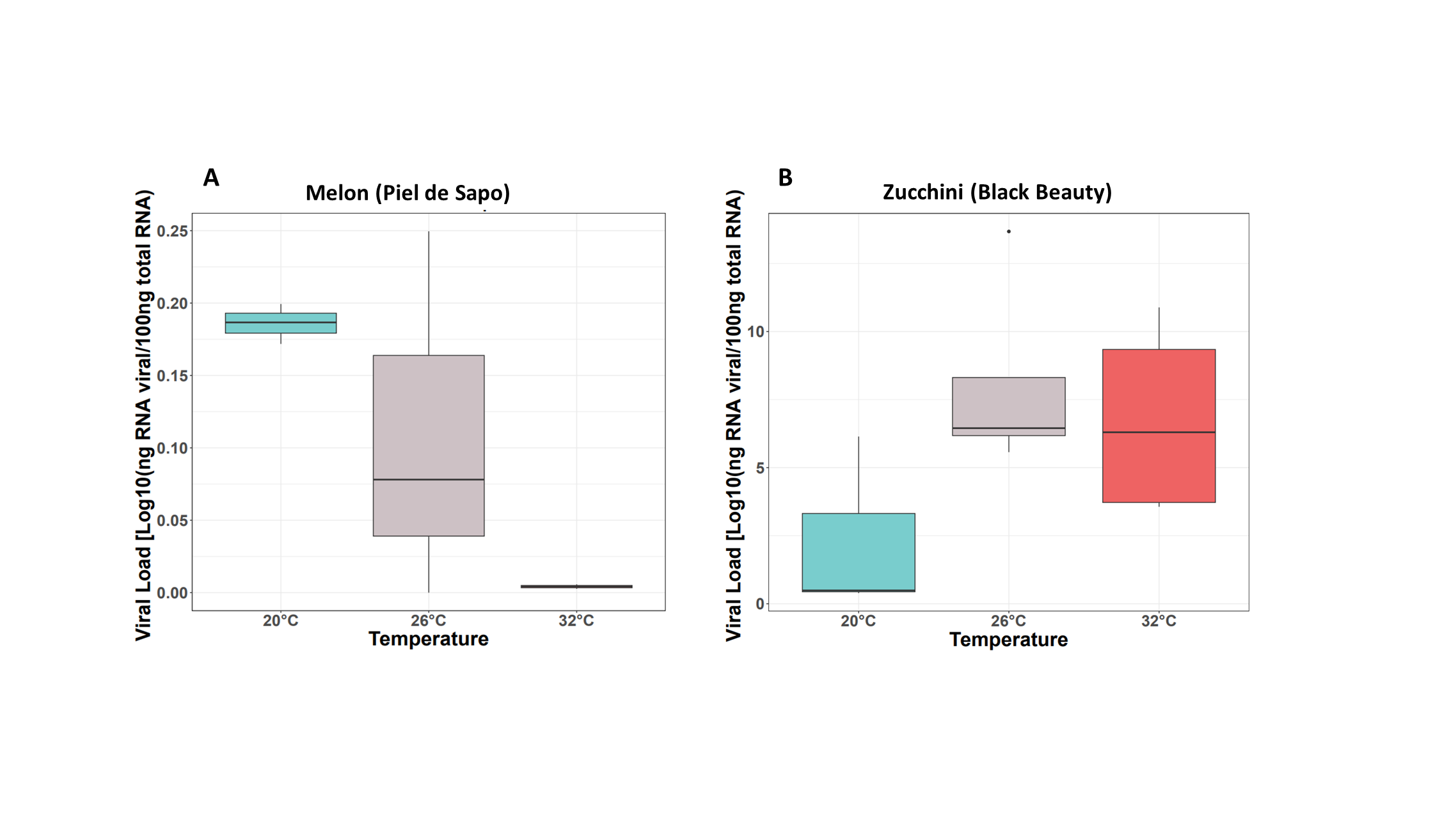

### Supp Figure 2

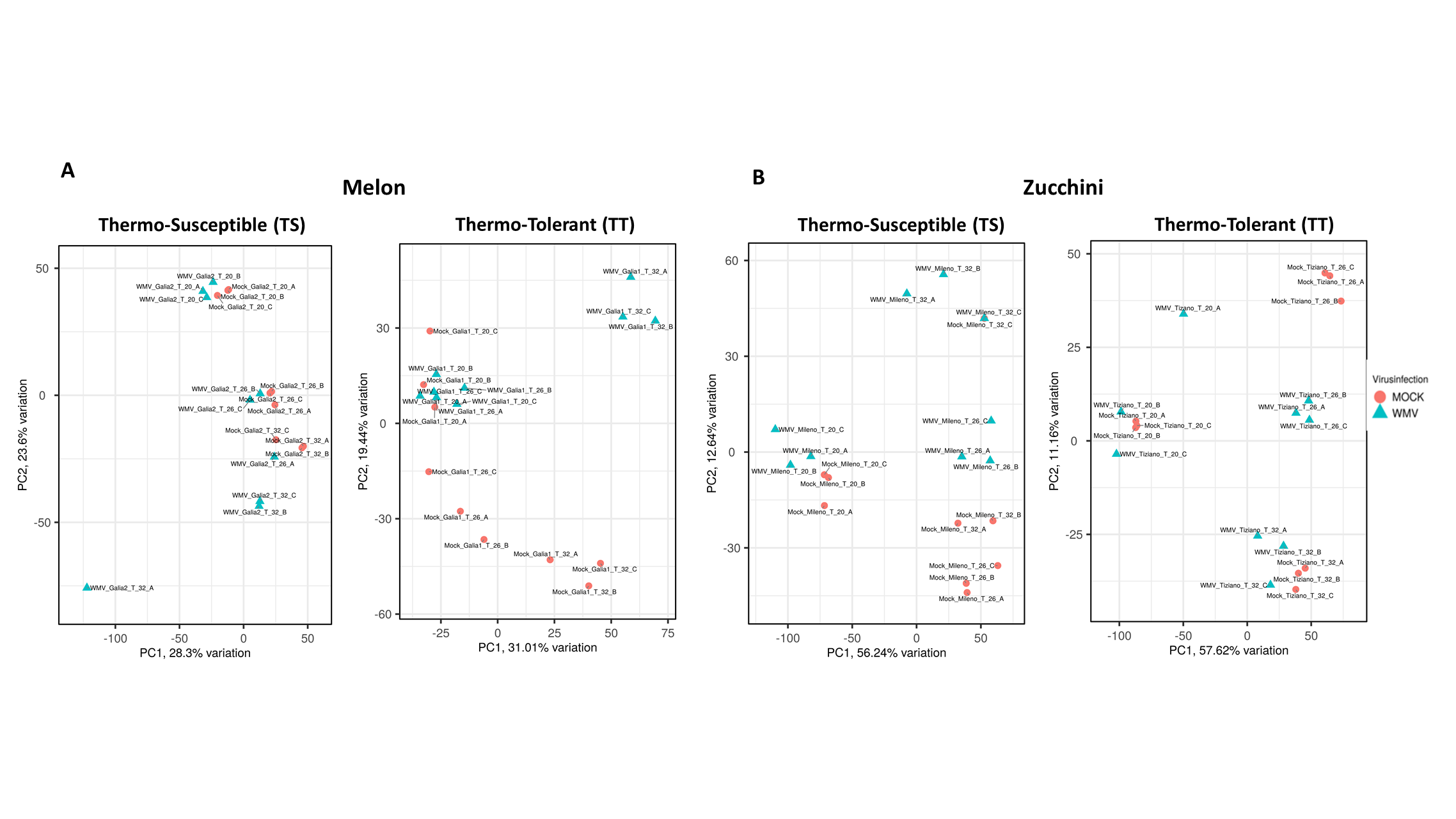

### Supp Figure 3

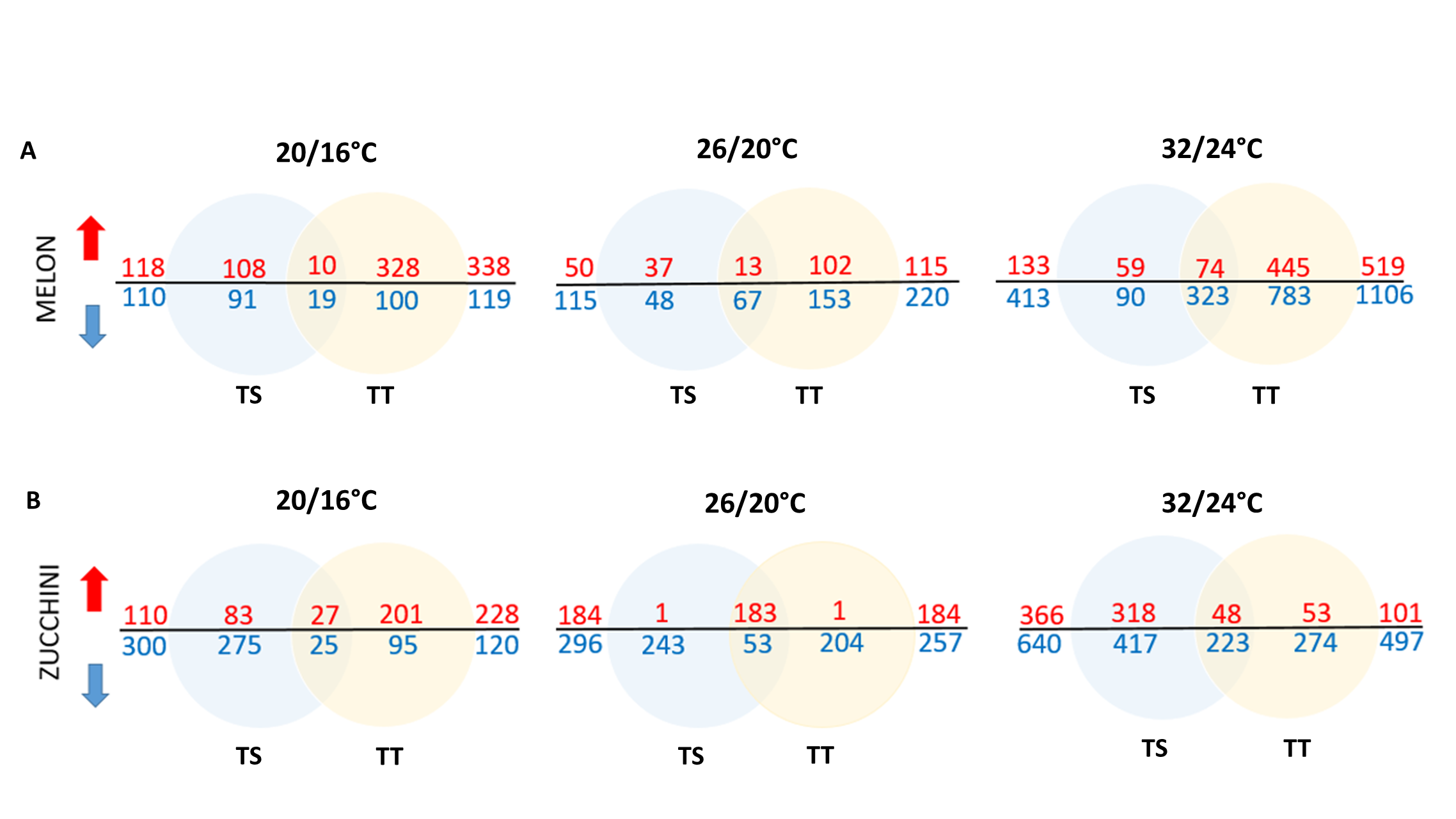
